# Heat-induced seizures, premature mortality, and hyperactivity in a novel *Scn1a* nonsense model for Dravet syndrome

**DOI:** 10.1101/2023.02.01.526678

**Authors:** Anat Mavashov, Marina Brusel, Jiaxing Liu, Victoria Woytowicz, Haneui Bae, Ying-Hsin Chen, Vardhan S. Dani, Elena Cardenal, Vittoria Spinosa, José Ángel Aibar, Moran Rubinstein

## Abstract

Dravet syndrome (Dravet) is a severe congenital developmental genetic epilepsy caused by *de novo* mutations in the *SCN1A* gene. Nonsense mutations are found in ~20% of the patients, and the R613X mutation was identified in multiple patients. Here we characterized the epileptic and non-epileptic comorbidities of a novel preclinical Dravet mouse model harboring this nonsense *Scn1a* mutation. Heterozygous *Scn1a* R613X mutation on a mixed C57BL/6J:129S1/SvImJ background exhibited spontaneous seizures, susceptibility to heat-induced seizures, and premature mortality, recapitulating the core epileptic phenotypes of Dravet. In addition, these mice, available as an open-access model, demonstrated increased locomotor activity in the open-field test, mimicking some non-epileptic Dravet-associated comorbidities. Conversely, *Scn1a*^WT/R613X^ mice on the pure 129S1/SvImJ background had a normal life span and were easy to breed. Homozygous *Scn1a*^R613X/R613X^ mice died before P16.

Our molecular analyses of hippocampal and cortical expression demonstrated that the premature stop codon induced by the R613X mutation reduced *Scn1a* mRNA and Na_v_1.1 protein levels to ~50% in heterozygous *Scn1a*^WT/R613X^ mice, with marginal expression in homozygous *Scn1a*^R613X/R613X^ mice. Together, we introduce a novel Dravet model carrying the R613X *Scn1a* nonsense mutation that can. be used to study the molecular and neuronal basis of Dravet, as well as the development of new therapies associated with *SCN1A* nonsense mutations in Dravet.

## Introduction

Dravet Syndrome (Dravet) is a severe childhood-onset developmental and epileptic encephalopathy. Most cases are caused by *de novo* mutations in the *SCN1A* gene, which encodes for the voltage-gated sodium channel Na_V_1.1 (Claes et al., 2003). Dravet patients develop normally in the first months of life. However, around six months of age, they start exhibiting severe febrile seizures, which soon progress to frequent spontaneous refractory seizures, episodes of status epilepticus, and a high incidence of sudden unexpected death in epilepsy (SUDEP). Additional non-epileptic comorbidities in Dravet include developmental delay and cognitive impairment (Dravet et al., 2011; Gataullina and Dulac, 2017; Cardenal-Muñoz et al., 2022).

Over 1,000 different *SCN1A* pathological variants have been identified in Dravet patients. Among these, ~7% are large deletions, 39% are missense mutations, and ~41% are nonsense or frameshift mutations (Claes et al., 2009; Meng et al., 2015; Xu et al., 2015; Liu et al., 2021). Most of these mutations are *de novo*, non-recurrent mutations. However, intriguingly, the *SCN1A* R613X nonsense mutation has been reported to occur *de novo* in ten different patients (Kearney et al., 2006; Margherita Mancardi et al., 2006; Depienne et al., 2009; Rodda et al., 2012; Wang et al., 2012; Gaily et al., 2013; Moehring et al., 2013; Lee et al., 2015). Here we set out to provide a comprehensive characterization of a novel open-access mouse model that harbors this nonsense mutation.

Dravet mouse models (DS mice) recapitulate many aspects of the human disease. To date, fifteen different models were developed, based on various *Scn1a* mutations. Importantly, all the DS mouse models recapture key pathophysiological phenotypes Dravet, exhibiting spontaneous seizures, premature mortality, and the presentation of Dravet-associated non-epileptic behavioral comorbidities (Yu et al., 2006; Ogiwara et al., 2007, 2013; Martin et al., 2010; Cheah et al., 2012; Miller et al., 2014; Tsai et al., 2015; Kuo et al., 2019; Ricobaraza et al., 2019; Dyment et al., 2020; Jansen et al., 2020; Uchino et al., 2021; Voskobiynyk et al., 2021; Valassina et al., 2022; Morey et al., 2022) (Table 1).

Several DS models are available through international repositories (Table 1), and two lines, developed by the Dravet Syndrome Foundation Spain, are distributed through the Jackson Laboratory: *i*) the conditional DS mice that harbor the *Scn1a*^A1783V^ missense mutation; and *ii*) a new *Scn1a*^R613X^ line, described here. While the first model has been validated and confirmed to recapitulate Dravet phenotypes (Table 1), the phenotypic and molecular characterization of the new *Scn1a*^R613X^ model was yet partial (Almog et al., 2022). Here, we show that DS mice carrying the *Scn1a*^R613X^ nonsense on a mixed C57BL/6J:129S1/SvImJ background recapitulate key phenotypes of Dravet, with heat-induced seizures that occur within the range of physiological fever temperatures at multiple developmental stages, spontaneous convulsive seizures, premature death, and hyperactivity in the open field. Moreover, these mice have reduced levels of *Scn1a* mRNA and Na_V_1.1 protein in the hippocampus and the cortex. Together, these data confirm that the *Scn1a*^R613X^ model is a valid animal model for Dravet research.

## Material and methods

### Animals

All animal experiments were approved by the Animal Care and Use Committee (IACUC) of Tel Aviv University. Mice used in this study were housed in a standard animal facility at the Goldschleger Eye Institute at a constant (22 °C) temperature, on a 12-h light / dark cycle, with *ad libitum* access to food and water.

Mice harboring the global *Scn1a^R613X^* mutation were generated by crossing males or females carrying an A-to-T point mutation in nucleotide 1837 (converting arginine 613 to a STOP codon) in addition to a silent C-to-T mutation at position 1833 (129S1/SvImJ-*Scn1a*^em1Dsf/J^, The Jackson Laboratory, stock no. 034129) (Fig. 1A), with wild-type (WT) mice, females or males (129S1/SvImJ, The Jackson Laboratory, stock no. 002448). Details about allele modification and genotyping are described here: https://www.jax.org/strain/034129. This mouse line was maintained on the pure 129S1/SvImJ genetic background. To produce DS mice, male or female *Scn1a^R613X^* mice on the pure 129S1/SvImJ were crossed with WT mice (males or females) on a C57BL/6J background (The Jackson Laboratory, stock no. 000664), generating F1 mice on a 50:50 genetic background. Both male and female offspring were used for experiments. Homozygous mutant mice (*Scn1a*^R613X/R613X^) were generated by crossing heterozygous *Scn1a*^WT/R613X^ mice on the pure 129S1/SvImJ background.

**Fig. 1:**
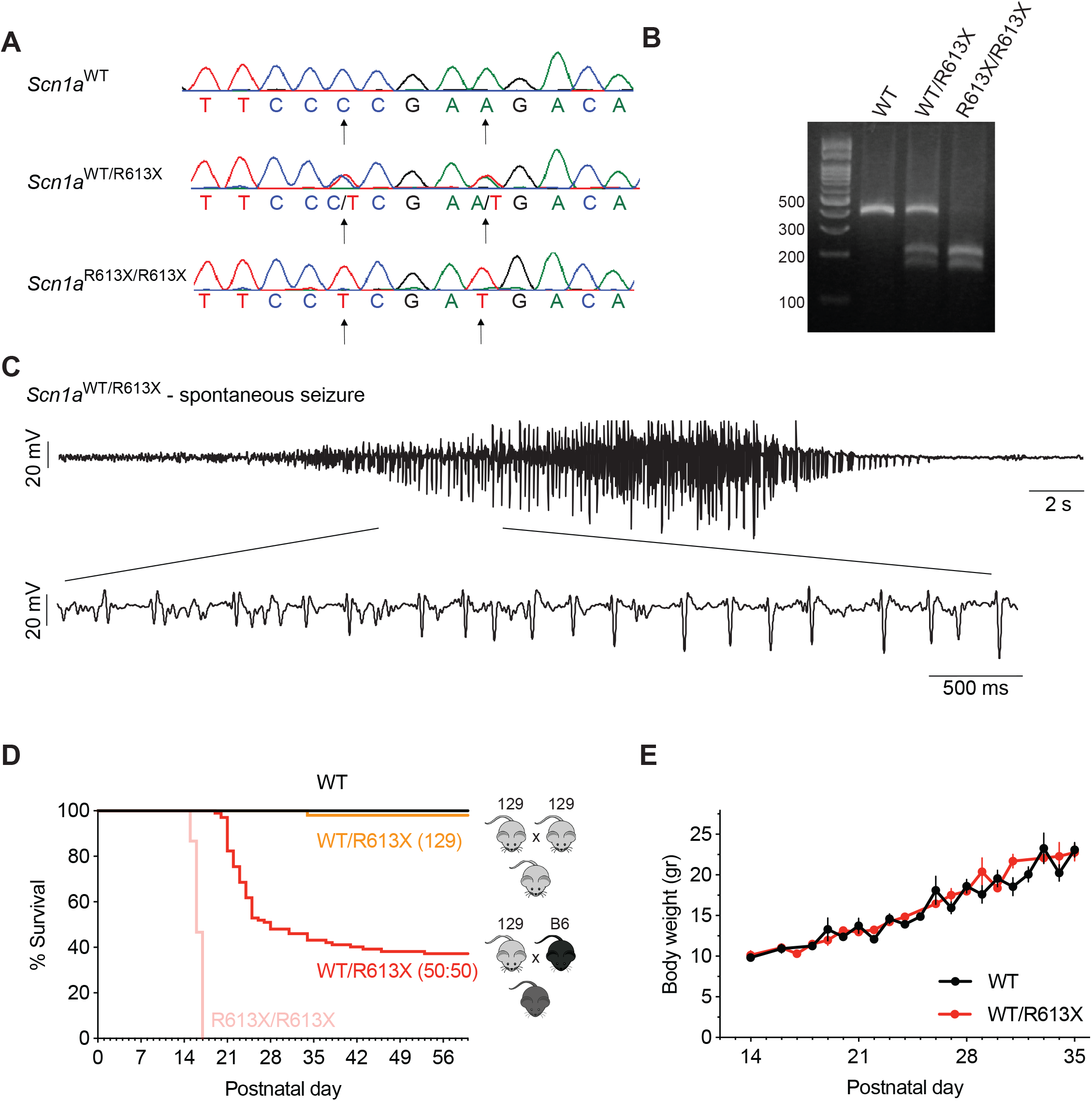
Spontaneous convulsive seizures and premature mortality in *Scn1a*^WT/R613X^ DS mice. **A.** CRISPR/Cas9-generated mutations were introduced to exon 12 of the mouse *Scn1a* gene: A>T point mutation at nucleotide 1837 (converting Arg613 into a STOP codon) and a silent C>T mutation at position 1833. **B.** Genotyping of *Scn1a*^R613X^ allele using PCR followed by *Taq*I digestion. **C.** A spontaneous seizure captured during an ECoG recording in a *Scn1a*^WT/R613X^ mouse on a mixed background. **D.** *Scn1a*^WT/R613X^ on a mixed background exhibited premature mortality, in contrast to *Scn1a*^WT/R613X^ mice on the pure 129S1/SvImJ. Homozygous *Scn1a*^R613X/R613X^ died prematurely between P14-16. (129 background: WT, n=24; *Scn1a*^WT/R613X^, n=51; *Scn1a*^R613X/R613X^, n=15. Mixed background: WT, n=30; *Scn1a*^WT/R613X^, n=102). **E.**The growth of *Scn1a*^WT/R613X^ mice on the mixed background was similar to their WT littermates. (WT, n=10; *Scn1a*^WT/R613X^, n=6).

### Genotyping

The PCR was performed using the primers and protocol described by the Jackson Laboratory (129S1/SvImJ-*Scn1a*^em1Dsf/J^, stock no. 034129). Following the amplification step, 7,000 units of *Taq*I restriction enzyme (New England Biolabs, Ipswich, MA, USA) were added to 6 μL of the PCR mixture, incubated at 65 °C for 15 minutes and analyzed on 3% agarose gel (Fig. 1B).

### Thermal induction of seizures

Thermal induction of seizures was performed as previously described (Almog et al., 2021). Briefly, the baseline body core temperature was recorded for at least 10 min, allowing the animals to habituate to the recording chamber and rectal probe. Body temperature was then increased by 0.5 °C every 2 min with a heat lamp (TCAT-2DF, Physitemp Instruments Inc., Clifton, NJ, USA) until a generalized tonic-clonic seizure was provoked; the temperature was not increased above 42 °C. Mice used for thermal induction of seizures were not included in the survival curve.

### Electrocorticography (ECoG) recordings

Electrode implantation was done at P21-P25, as previously described (Fadila et al., 2020). Mice were allowed to recover for at least 48 hours before recording.

### Behavioral experiments

Behavioral tests were done as described previously (Fadila et al., 2020). For the open-field test, the mice were placed in the center of a square (50 × 50 cm) plexiglas apparatus and their movement was recorded for 10 min. Live tracking was achieved via a monochrome camera (Basler acA1300-60gm, Basler AG, Ahren, Germany) connected with EthoVision XT 13 software (Noldus Technology, Wageningen, Netherlands). For analysis of anxiety-like behavior, the software was set to subdivide the open field into 10 × 10 cm squares (25 in total). Those squares immediately adjacent to the apparatus walls constituted an “outer” zone, while the remaining squares not touching the walls demarcated an “inner” zone.

To examine motor functions, the mice were placed on an accelerating rotating rod (acceleration from 3 to 32 RMP, Med Associates, Inc., VT, USA), and the time at which each mouse fell was recorded. The test was repeated five times for each mouse, and the three longest trials were averaged.

To test spatial working memory, a symmetrical Y-maze comprised of three arms (each 35 cm L × 7.6 cm W × 20 cm H) was used. The mouse was placed into one of the Y-maze arms and allowed free exploration for 10 min. Live tracking was achieved via a monochrome camera (Basler acA1300-60gm, Basler AG, Ahren, Germany) connected with EthoVision XT 13 software (Noldus Technology, Wageningen, Netherlands). Different cohorts of mice were tested at each time point to ensure a reaction to a novel arena. The analysis included the number of arms entered, excluding the first recorded arm in which the mouse was placed. The sequence of entries was recorded, as well as the number of triads (entries into a set of 3 different arms, in any sequence). The percentage of spontaneous alternation was calculated as the number of triads divided by the number of possible triads.

### qPCR

Total RNA was isolated using Purelink^™^ RNA mini kit according to the manufacturer’s instructions (Thermo Fisher Scientific, Life Technologies, Carlsbald, CA, USA). cDNA was synthesized from 500 ng RNA using Maxima H Minus cDNA synthesis kit (Thermo Fisher Scientific, Life Technologies, Carlsbald, CA, USA). Real-time PCR (qPCR) reactions were performed in triplicates in a final volume of 10 μL with 5 ng of RNA as template using the TaqMan *Scn1a (*Mm00450580_m1) gene expression assay (Applied Biosystems, Thermo Fisher Scientific, Life Technologies, Carlsbald, CA, USA). Two endogenous controls were used: *Gusb (*Mm00446953_m1) and *Tfrc (*Mm00441941_m1). Efficiency of 100%, dynamic range, and lack of genomic DNA amplification were verified for all the assays.

### Western blot

The hippocampi were extracted and homogenized as previously described (Nissenkorn et al., 2019). Briefly, 0.45-0.7 mg of tissue was homogenized in 0.32 M sucrose supplemented with protease inhibitors (Sigma-Aldrich, St. Louis, MO, USA), 1mM EDTA and 1 mM PMSF, pH 7.4. Crude membrane preparation was produced by centrifugation at 17,000 × g for 75 min, as previously described (Nissenkorn et al., 2019). The pellet was solubilized in 150 mM NaCl, 2% Triton X-100, 25 mM Tris, supplemented with protease inhibitors, 1 mM EDTA and 1 mM PMSF, pH 7.4. 50 μg aliquots of total protein were separated on Tris-acetate gel (6%) and transferred onto PVDF membrane. After overnight blocking in 5% nonfat dry milk in Tris-buffered saline (TBS), the membrane was incubated overnight with anti-Na_V_1.1 antibody (1:200, Alomone Labs, Jerusalem, Israel; Catalog# ASC-001) or anti-calnexin (1:2,000, Stressgen Biotechnologies, San Diego, USA), followed by 2 h incubation with HRP-conjugated goat anti-rabbit antibody (1:10,000, Sigma-Aldrich, St. Louis, MO, USA). The signal was visualized by chemiluminescent detection using ECL.

### Digital PCR (dPCR) for allele specific quantification of WT and R613X mRNA

Allele-specific dPCR assays were performed at Tevard Biosciences (Cambridge, MA, USA) on neocortical tissue isolated from WT and *Scn1a*^WT/R613X^ mice on a mixed 50:50 C57BL/6J: 129S1/SvImJ background, generated using a similar breeding strategy as described above. After confirming anesthesia, neocortical tissue was carefully extracted, flash-frozen in liquid nitrogen, and stored at −70 °C until processing. Total RNA was extracted from tissues using AllPrep^®^ DNA/RNA/miRNA Universal kit (Qiagen, Hilden, Germany; Catalog# 80224) and reverse transcribed with SuperScript^™^ IV First-Strand Synthesis System (Thermo Fisher, Waltham, MA, USA; Catalog# 18091050). For allele-specific detection of *Scn1a* transcripts (WT vs. R613X mRNA), custom-designed Affinity Plus^™^ qPCR Probes (IDT, Coralville, IA, USA) were used to enable greater SNP target specificity. The sequences for the primers and probes are as follows, where the + sign before a nucleotide indicates locked nucleotides: Forward primer: CACAGCACCTTTGAGGATAAT; Reverse primer: GGTCTGGCTCAGGTTACT; WT probe: TCC+C+CG+A+A+GAC; R613X probe: TCC+T+CG+A+T+GAC+AC. Mouse Gapdh TaqMan probe (Thermo Fisher; Mm99999915_g1) was used for quantification of *Gapdh* as a reference gene. Digital PCR (dPCR) for absolute quantification of gene targets was performed using naica^®^ System dPCR from Stilla Technologies (Villejuif, France).

### Meso Scale Discovery Electrochemiluminescence (MSD-ECL) assay

MSD-ECL assays were performed at Tevard Biosciences (Cambridge, MA, USA). Total protein was isolated from combined neocortex (parietal, temporal, occipital lobes), or dissected cortical lobes (frontal, parietal, temporal, occipital, respectively) and liver. The tissue was homogenized in lysis buffer (1x TBS, 1% TX-100, 0.5% Nonidet P-40, 0.25% Na deoxylate, 1mM EDTA) supplemented with protease/phosphatase inhibitors (HALT Protease/Phosphatase inhibitor cocktail, Thermo Fisher Catalog # 78440) using beaded tubes (MP Biomedicals, Irvine, CA, USA Catalog # 116913050-CF) and a benchtop homogenizer (MP Biomedicals FastPrep-24 5G) at 4.0 m/s for 5 seconds. Samples were incubated at 4 °C for 15 minutes with rotation, spun down at 16000 × g at 4 °C for 15 minutes, and the supernatant containing total protein was collected. Multi-Array 96 Small Spot Plates (Meso Scale Diagnostics, Rockville, MD, USA, Catalog # L45MA,) were blocked for 1 hour at room temperature with shaking using Blocker B (Meso Scale Diagnostics, Catalog # R93BB). Plates were washed 3 times with TBST, then coated with 5 μg/mL capture antibody for Na_V_1.1 (UC Davis, Antibodies Inc. Catalog # 75-023) at room temperature for 1 hour with shaking. Plates were washed as previously described, and 25 μL of 4 mg/mL standard or protein samples were added to individual wells and incubated overnight at 4 °C with shaking. Plates were washed, and the detection antibody for Na_V_1.1 (Alomone Labs, Catalog # ASC-001-SO) was added at 2.4 μg/mL and incubated at RT for 1 hour with shaking. Plates were washed, and secondary detection antibody (SULFO-TAG anti-rabbit antibody (Meso Scale Diagnostics Catalog# R32AB) was added at 2 μg/mL and incubated for 1 hour at room temperature with shaking. Plates were washed and read using MSD GOLD Read Buffer B (Meso Scale Diagnostics Catalog # R60AM, 150 μL) on the MSD QuickPlex SQ120 Instrument.

## Results

### DS *Scn1a*^R163X^ mice on the mixed background demonstrated premature mortality and spontaneous seizures

The *Scn1a*^R613X^ mutation was generated on the 129S1/SvImJ background (Fig. 1A, B). Heterozygous *Scn1a*^WT/R613X^ mice on the pure 129S1/SvImJ background did not exhibit spontaneous convulsive seizures, as observed during routine handling, and only one mouse died prematurely, in agreement with previous reports (Yu et al., 2006; Miller et al., 2014). Conversely, crossing these mice onto the C57BL/6J background uncovered their epileptic phenotypes. *Scn1a*^WT/R613X^ on a mixed background (50:50 C57BL/6J: 129S1/SvImJ) exhibited spontaneous convulsive seizures, which were often observed during routine handling (Fig. 1C, D and Video. S1). Moreover, these mice demonstrated profound premature mortality, with only ~40% of mice surviving to P60, and most of the deaths occurring during their fourth week of life (P21-P28, Fig. 1D). Despite poor survival, the growth rate of *Scn1a*^WT/R613X^ mice on the mixed background was similar to their WT littermates, with no significant differences in body weight (Fig. 1E). Homozygous *Scn1a*^R613X/R613X^ died prematurely between P14-16 (Fig. 1D).

Together, the *Scn1a*^WT/R613X^ Dravet mice on a mixed background demonstrated spontaneous seizures and premature mortality, recapitulating Dravet epilepsy and Dravet-associated premature death.

### DS *Scn1a*^R163X^ mice exhibit heat-induced seizures and hyperactivity in the open field

Heat-induced seizures are a hallmark phenotype of *SCN1A* mutations. We tested the susceptibility of *Scn1a*^WT/R613X^ mice, on either background, to heat-induced seizures at different ages: the pre-epileptic stage (P14-P16), the severe stage of epilepsy during the fourth week of life, and the stabilization or chronic stage after the fifth week of life (Fadila et al., 2020; Gerbatin et al., 2022). All the *Scn1a* mutant mice exhibited thermally induced seizures below 42°C (Fig. 2). The highest susceptibility, as demonstrated by the lowest temperature threshold, was observed in *Scn1a*^WT/R613X^ on the mixed background at their fourth week of life (P21-P25, Fig. 2B-C, F). Together, these data confirm that *Scn1a*^WT/R613X^ mice show a high susceptibility to heat-induced seizures, from the pre-epileptic stages to adulthood, similar to Dravet patients.

**Fig. 2.**
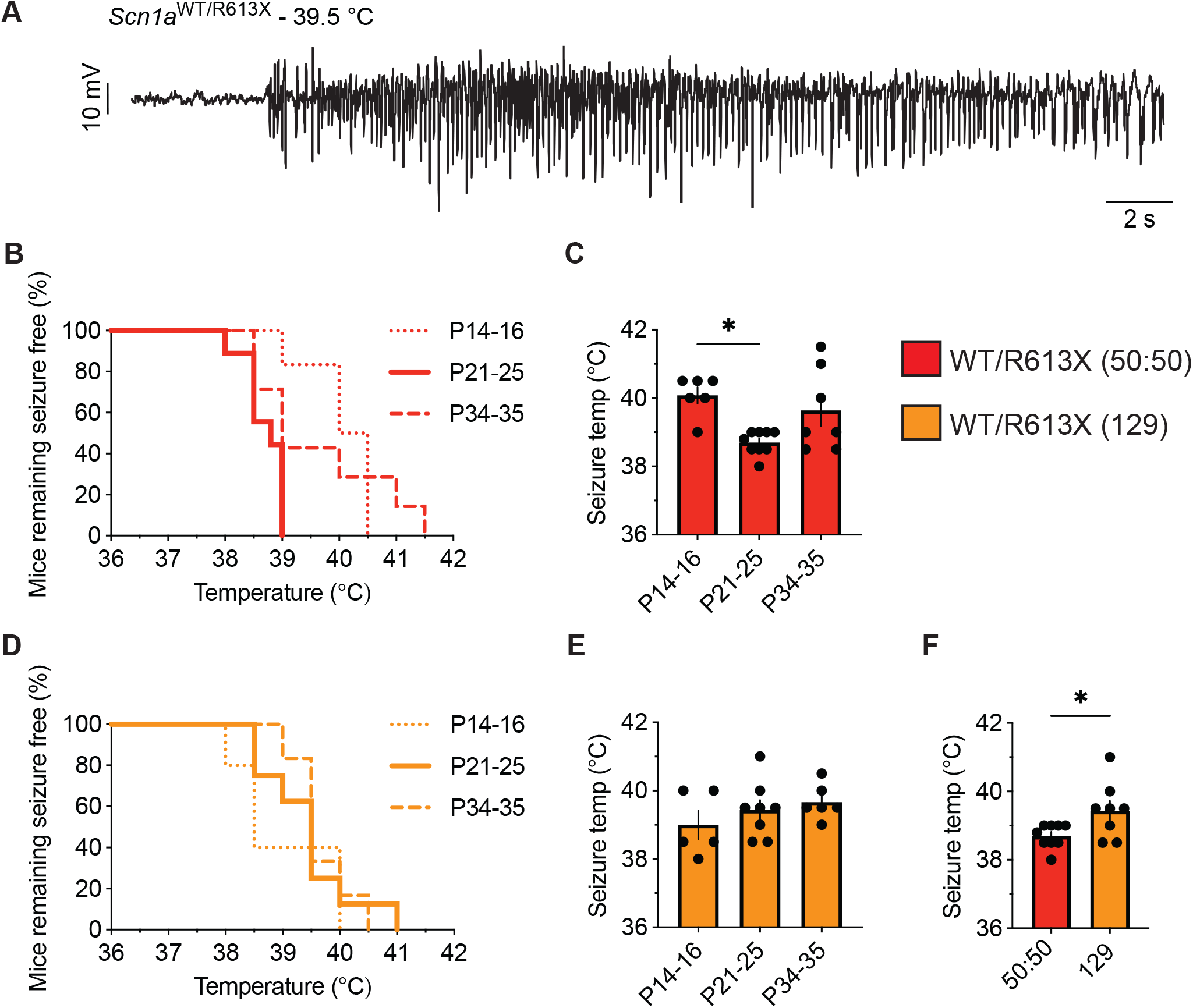
*Scn1a*^WT/R613X^ are susceptible to heat-induced seizures. **A.** An EcoG trace from a *Scn1a*^WT/R613X^ mouse on a mixed 50:50 background depicting epileptic activity at a temperature of 39.5 °C. **B-C.** *Scn1a*^WT/R613X^ mice on a mixed background, at the indicated ages, remaining free of thermally induced seizures (**B**), and the temperature of seizures (**C**). P14-16: n=6; P21-P25: n=9; P34-P35: n=7. **D-E.** *Scn1a*^WT/R613X^ mice on the 129S1/SvImJ background, at the indicated ages, remaining free of thermally induced seizures (**D**), and the temperature of seizures (**E**). P14-16: n=5; P21-P25: n=8; P34-P35: n=6. WT mice did not experience seizures within this temperature range (n=4-7 at each age group, not shown). **F**. At P21-P25, *Scn1a*^WT/R613X^ mice on a mixed background have heat-induced seizures at lower temperatures compared to *Scn1a*^WT/R613X^ mice on a pure 129 background. These are the same data depicted in C and E. These data did not follow normal distribution. Statistical comparison in C utilized nonparametric One-Way ANOVA followed by Dunn’s test. Statistical comparison in E utilized the Mann-Whitney test. *, p<0.05

To test the presence of non-epileptic comorbidities, we examined hyperactivity in a novel arena, motor functions using the rotarod test, and working memory using the WT/R613XY-maze spontaneous alternation test in *Scn1a*^WT/R613X^ mice on the mixed background. In the open field test, which examines the exploration of a novel arena, heterozygous *Scn1a*^WT/R613X^ mice displayed increased ambulation and traveled longer distances and at higher velocities compared to their WT littermates (Fig. 3A-C). Nevertheless, the time in the center of the arena was similar between *Scn1a*^WT/R613X^ and WT mice (Fig. 3D), indicating that these mice do not exhibit increased anxiety. Next, motor functions were assessed using the rotarod test. As shown in Fig. 3E, the latency to fall was comparable between WT and heterozygous *Scn1a*^WT/R613X^ mice, suggesting normal balance and coordination. Moreover, their performance in the spontaneous alternation Y-maze during their fourth week of life, corresponding to the severe stage of Dravet (P21-P25), as well as in a different cohort of mice at their sixth week of life (P36-P43), indicated that these mice do not have a deficit in spatial working memory (Fig. 3F, H). Nevertheless, increased locomotor activity was also observed in the Y-maze, further corroborating the hyperactivity observed in the open field. Thus, in addition to susceptibility to thermally induced seizures, spontaneous seizures, and premature death, *Scn1a*^WT/R613X^ mice on the mixed background also demonstrate hyperactivity.

**Fig. 3.**
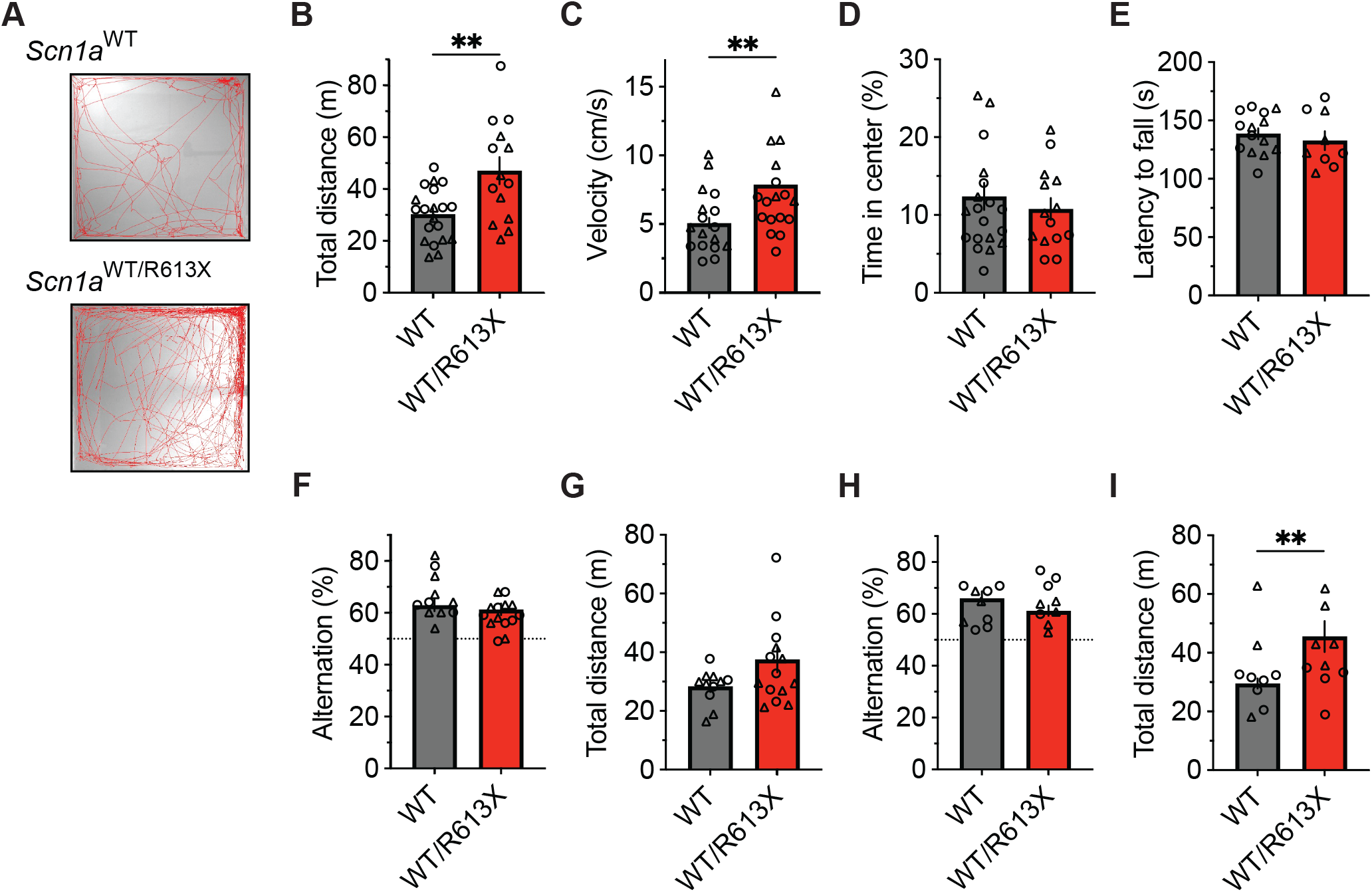
Hyperactivity in *Scn1a*^*WT*/R613X^ mice on the mixed background. **A-D.** Open field in WT and DS mice at their fourth week of life (P21-P25). **A.** Representative examples of exploring the behavior of WT and DS mice during the 10-minute test period. **B**. Distance moved. **C**. Average velocity. **D.** Percentage of time spent in the central portion of the arena. WT, n=20; *Scn1a*^WT/R613X^, n=14. **E.** The latency to fall on the rotarod test in WT and DS mice (P34-P42). WT, n=14; *Scn1a*^WT/R613X^, n=9. **F-I**. The Y-maze in P21-P25 mice (**F-G**) and in another cohort of older (P36-P43) mice (**H-I**). The dotted line in F and H signifies the chance level expected from random alternation. G and I depict the total distance moved in the Y-maze during exploration. P21-P25: WT, n=14; *Scn1a*^WT/R613X^, n=11; P36-P43: WT, n=10; *Scn1a*^WT/R613X^, n=8. In addition to the average and SE, the individual data points from males (triangles) and females (circles) are depicted. As these data were distributed normally, statistical comparison unpaired t-test. **, p<0.01

As *Scn1a*^WT/R613X^ mice on the pure 129S1/SvImJ background did not exhibit premature mortality (Fig. 1) but were susceptible to heat-induced seizures (Fig. 2), we wondered if these mice exhibited behavioral deficits. While we did find vast differences in the innate tendency to explore the novel arena (compare Figs. 3 and 4), the activity of *Scn1a*^WT/R613X^ mice on the 129S1/SvImJ background was similar to that of their WT littermate in both the open field test and the rotarod test, suggesting that genetic background can also modify Dravet-associated hyperactivity.

**Fig. 4.**
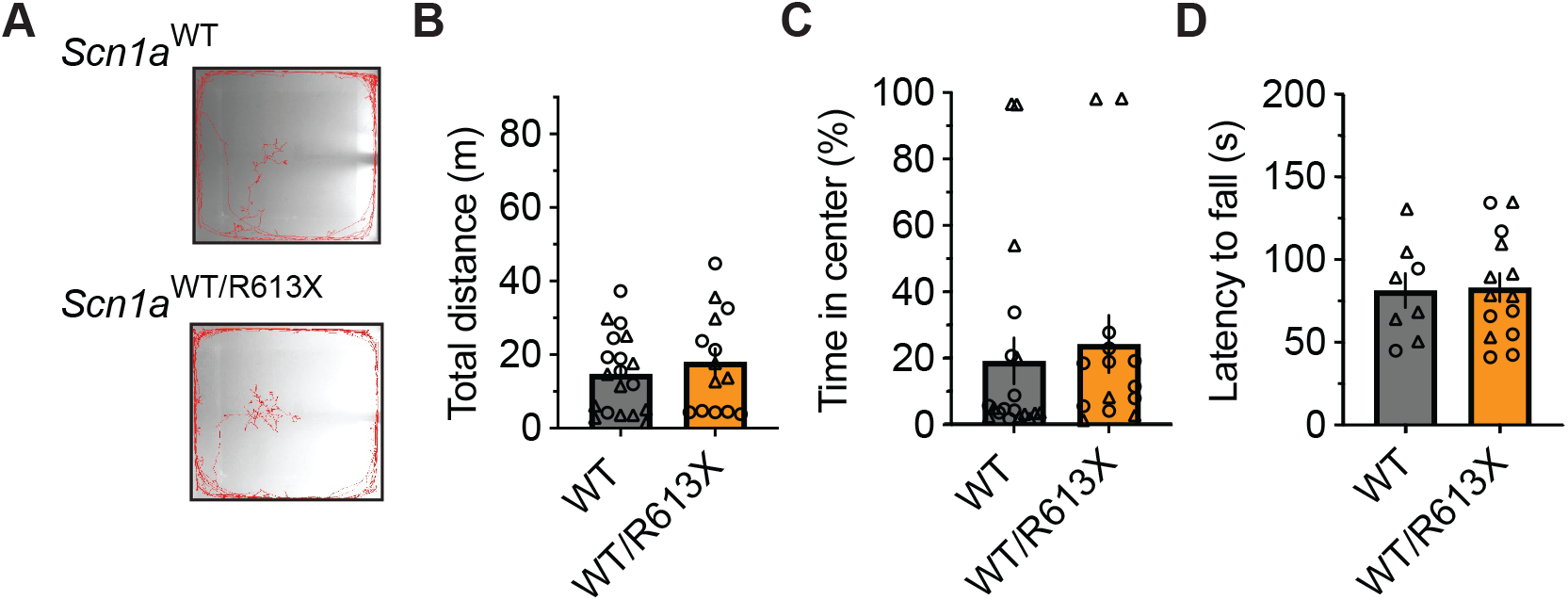
No behavioral deficits in *Scn1a*^*WT*/R613X^ mice on the 129S1/SvImJ background. **A.** Representative examples of exploring the behavior of wild-type (WT) and *Scn1a*^WT/R613X^ mice during the 10-minute test period. **B**. Distance moved in the open field. **C**. Percentage of time spent in the central portion of the arena. WT, n=19; *Scn1a*^WT/R613X^, n=14. **D.**The latency to fall on the rotarod test. WT, n=8; *Scn1a*^WT/R613X^, n=14.

### Reduced *Scn1a* mRNA and protein expression

To test the effect of the R613X premature termination at the transcriptional level, we extracted and analyzed mRNA from the hippocampus of P21-P24 WT and *Scn1a* mutant mice. First, we confirmed that the R613X mutation is also expressed in the hippocampus and translated into mRNA (Fig. 5A). Next, quantitative real-time PCR analysis of *Scn1a* mRNA, using an assay that targets the boundary junction between exons 18-19, demonstrated ~50% reduction in expression of the full-length *Scn1a* mRNA in heterozygous *Scn1a*^WT/R613X^ mice on either genetic background and a marginal background expression in homozygous *Scn1a*^R613X/R613X^ mice (Fig. 5B, C).

**Fig. 5.**
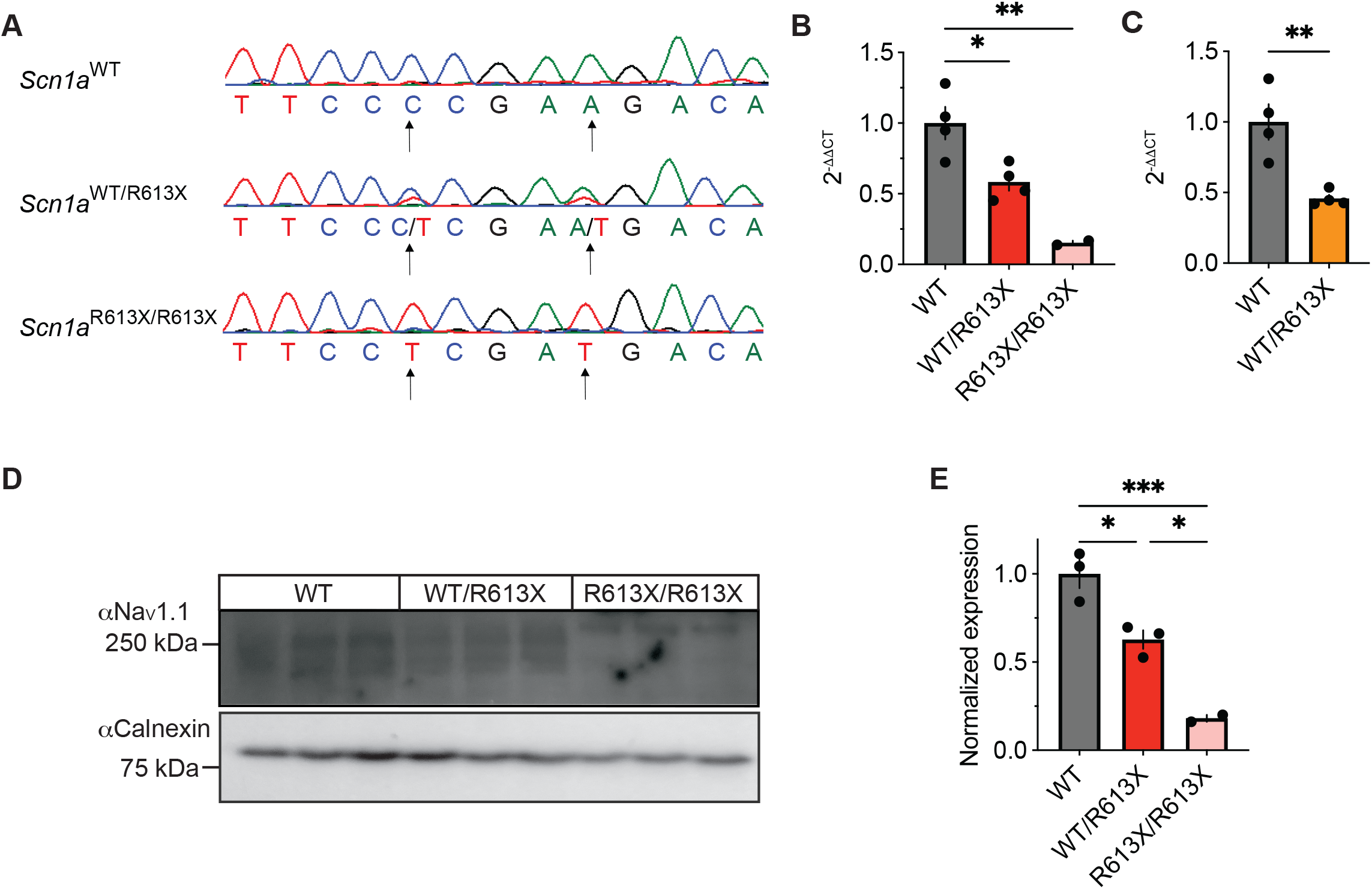
Reduced *Scn1a* mRNA and Na_V_1.1 protein expression in the hippocampus of *Scn1a^WT/^*^R613X^ mice. **A**. Sequencing of hippocampal mRNA confirmed the expression of the *Scn1a*^R613X^ allele. **B-C.** qPCR analysis of *Scn1a* mRNA expression levels showed a significant reduction to approximately 50% in heterozygous *Scn1a*^WT/R613X^ on the mixed background (B), as well as on the pure 129S1/SvImJ background (C). A marginal background expression was found in homozygous *Scn1a*^R613X/R613X^ mice. Mixed background: WT, n=4; *Scn1a*^WT/R613X T^, n=4; *Scn1a*^R613X/R613X^, n=2. 129S1/SvImJ background: WT, n=4; *Scn1a*^WT/R613X^, n=4. **D.** Western blot analysis of Na_V_1.1 protein extracted from the hippocampi of three different mice from each genotype. **E**. Quantification of Na_V_1.1 protein level from D. Only two of the three *Scn1a*^R613X/R613X^ were included in the quantification due to the dark spot in the middle lane that precluded the inclusion of this sample. Statistical comparison in C and E utilized One-Way ANOVA followed by Tukey’s multiple comparisons test. Statistical comparison in C utilized t-test. *, p<0.05, **, p<0.01, ***, p<0.001.

Next, we determine the impact of the R613X mutation on Na_V_1.1 protein expression levels. In accordance with the decrease observed in *Scn1a* mRNA, western blot analysis of extracted hippocampi showed that the level of Na_V_1.1 protein in the hippocampus of heterozygous *Scn1a*^WT/R613X^ was decreased to ~50% compared to WT control, with a minimal signal seen in homozygous *Scn1a*^R613X/R613X^ mice (Fig. 5D, E).

To quantitatively assess the level of the R613X transcript in the cortex, we performed an allele-specific dPCR assay, using probes that specifically bind WT *Scn1a* and R613X sequences (Fig. 1A). The R613X allele was not detected in cortical tissue from WT mice, confirming the specificity of the assay (Fig. 6A). Conversely, the WT allele was detected at about half the levels in *Scn1a*^WT/R613X^ tissue compared to WT cortex, as expected for heterozygous animals (Fig. 6B). Interestingly, when expressed as the percentage of the WT allele, the steady-state level of the mutant R613X transcript was 8.9 ±0.9% (Fig. 6C), much lower than the expected 50% level, suggesting that transcripts of the mutant allele undergo strong nonsense-mediated decay (NMD) (Jaffrey and Wilkinson, 2018) in neocortical tissue.

**Fig. 6.**
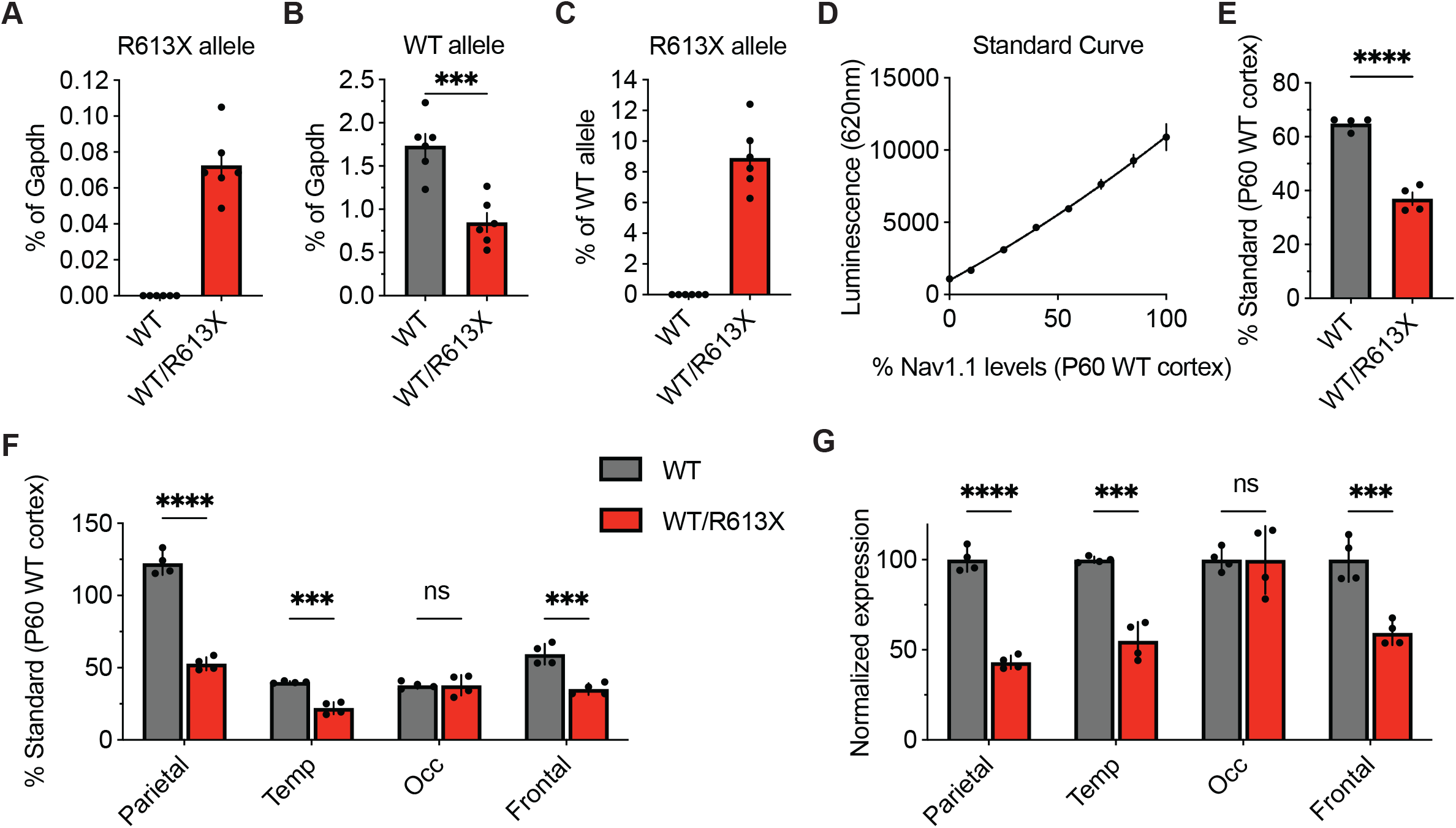
Reduced *Scn1a* mRNA and Na_V_1.1 protein expression in the cortex of *Scn1a^WT/R613X^* mice. **A-C.** Allele-specific quantification of *Scn1a* transcripts via dPCR. The R613X and WT *Scn1a* alleles were quantified with Affinity Plus probes on dPCR and normalized to mouse *Gapdh* levels. **A.**Cortical tissue from WT mice showed no detectable levels of the R613X allele transcript, demonstrating the specificity of allelic discrimination using this assay. **B.**In cortical tissue from *Scn1a*^WT/R613X^ mice, the WT *Scn1a* allele is at 48.8% compared to in WT cortical tissue. **C.** Shown as the ratio of R613X to WT alleles, the mean steady-state level of the *Scn1a* R613X allele is at 8.9±0.9% of the WT allele in *Scn1a*^WT/R613X^ animals. WT, n=6; *Scn1a*^WT/R613X^, n=6. **D-F.** Cortical levels of Na_V_1.1 proteins were quantified using the Meso Scale Discovery Electrochemiluminescence (MSD-ECL) assay. **D.** The standard curve was generated by mixing P60 WT cortical and liver proteins at different ratios. The resulting 2^nd^ order polynomial fit, depicted as a solid line (R^2^ = 0.9924) was used to calculate protein levels relative to P60 WT cortex in E and F. **E.**Protein Na_V_1.1 expression in the whole cortex. **F.** Na_V_1.1 protein expression in the parietal, temporal, frontal, and occipital lobes of the neocortex of WT and WT/R613X mice. **G.** The same data from F, but normalized to Na_V_1.1 expression in WT mice. WT, n=4; *Scn1a*^WT/R613X^, n=4. Statistical comparison by unpaired t-test. ***, p<0.001, ****, p<0.0001

For quantitative analysis of Na_V_1.1 protein levels, we performed the Meso Scale Discovery Electrochemiluminescence (MSD-ECL) assay. MSD-ECL is a highly sensitive ELISA-based assay that enables the detection of small changes in protein levels. Here, we quantified Na_V_1.1 protein levels in four different cortical lobes (parietal, temporal, occipital, and frontal) on MSD-ECL utilizing two distinct anti-Na_V_1.1 antibodies (Fig. 6D-F). Overall, in the cortex of *Scn1a*^WT/R613X^, the level of NaV1.1 was reduced to about 50% compared to WT mice (Fig. 6E). However, we found variation in the level of Na_V_1.1 across the cortex, with the highest Na_V_1.1 expression in the parietal lobe, and lower levels in other regions (Fig. 6F). NaV1.1 expression was reduced to approximately 50% in the parietal, temporal and frontal, but similar to that of WT mice in the occipital lobe (Fig. 6G). Together, these data confirm reduced *Scn1a* mRNA and Na_V_1.1 protein expression in *Scn1a*^WT/R613X^ mice, in both the hippocampus and the cortex.

## Discussion

Dravet is a severe form of developmental and epileptic encephalopathy with limited treatment options and poor prognosis. To date, fifteen different models have been generated based on microdeletions, nonsense, or missense mutations in the *Scn1a* gene (Table 1). The novel *Scn1a*^WT/R613X^ model described here demonstrated core Dravet-associated phenotypes which include spontaneous convulsive seizures, high susceptibility to heat-induced seizures, premature mortality, and several nonepileptic behavioral comorbidities. Thus, this model is open-access and publicly available model can be used by the Dravet community for preclinical studies of Dravet mechanisms and therapies.

Genetic background was shown to dramatically modulate the effect of *Scn1a* mutations dramatically. DS mice on the pure C57BL/6J background demonstrate the most severe phenotypes, with high mortality of 60-80% and frequent spontaneous seizures. Conversely, mice with the same *Scn1a* mutation on the pure 129X1/SvJ or 129S6/SvEvTac background rarely experience spontaneous seizures or premature death (Yu et al., 2006; Miller et al., 2014; Mistry et al., 2014; Rubinstein et al., 2015; Kang et al., 2018). In accordance, *Scn1a*^WT/R613X^ mice on the pure 129S1/SvImJ background have a normal life span (Fig. 1) and unaltered behavior in the open field test (Fig. 4). Despite that, we did observe susceptibility to heat-induced seizures (Fig. 2), as well as reduced expression of *Scn1a* mRNA in the hippocampus (Fig. 5).

Multiple studies have used DS mice models to examine the therapeutic potential of current and novel drug treatments (Hawkins et al., 2017; Han et al., 2020; Isom and Knupp, 2021; Pernici et al., 2021; Tanenhaus et al., 2022). Our characterization highlights several readouts that may be useful in future studies to examine the therapeutic benefit while using *Scn1a*^WT/R613X^ mice on a mixed C57BL/6J: 129S1/SvImJ background. key Dravet-associated phenotypes in these mice include: *i*) spontaneous convulsive seizures (Fig. 1C and Video S1); *ii*) profound premature mortality (Fig. 1D) with overall survival of less than 50%; *iii*) high susceptibility to thermally-induced seizures (Fig. 2) with heat-induced seizures at multiple developmental stages occurring within the range of physiological fever temperatures. Heat-induced seizures at relatively low temperatures provide an advantage with a wide measurement range to quantify the effect of current and novel treatments, with the highest sensitivity around P21 (Fig. 2C); *iv*) developmental changes in the severity of the epileptic phenotypes. Specifically, the pre-epileptic stage in *Scn1a*^WT/R613X^ DS mice was characterized by susceptibility to heat-induced seizures that preceded the onset of premature mortality (P14-P16); profound mortality during the fourth week of life, with 78% of death occurring between P21 and P28, corresponding to the severe or worsening stage of Dravet, and some stabilization with a reduced rate of premature death in mice that survive beyond P30 (Fig. 1D). Thus, relatively restricted age groups should be considered for analysis; *v*) the presentation of non-epileptic comorbidities demonstrated here as motor hyperactivity when introduced to a novel arena (Fig. 3), modeling Dravet-associated hyperactivity. Of note, was also observed hyperactivity in the Y-maze test (Fig. 3), indicating that this is a robust and reliable readout for Dravet-associated nonepileptic phenotype; *vi*) reduced *Scn1a* mRNA and Na_V_1.1 protein levels (Figs. 5, 6); *vii*) impaired firing of inhibitory neurons, a typical neuronal deficit in DS mice, was also observed previously in *Scn1a*^WT/R613X^ mice (Almog et al., 2022). Importantly, these robust phenotypes highlight this model as an open-access pre-clinical platform to study Dravet therapies.

Nevertheless, some of the non-epileptic phenotypes observed in other DS models were not detected here (Table 1), possibly due to the effect of genetic background on the presentation of Dravet-associated non-epileptic comorbidities. Specifically, while *Scn1a*^WT/R613X^ mice demonstrated motor hyperactivity when introduced to a novel arena (Fig. 3A-C), no motor deficits and altered spatial working memory were observed using the rotarod or the Y-maze spontaneous alternation tests (Fig. 3). However, these data are in accordance with other DS models that were reported to have normal rotarod performance, including models with deletions in the *Scn1a* gene (Niibori et al., 2020; Patra et al., 2020; Valassina et al., 2022) or mice harboring the *Scn1a* R1407X nonsense mutation (Ito et al., 2013). Similarly, normal spontaneous alternation were also reported in DS mice with deletion of the 1^st^ exon or mice with knock in of Scn1a poison exon (Voskobiynyk et al., 2021; Gerbatin et al., 2022).

Conversely, impaired rotarod activity and spontaneous alternation was reported in DS mice with deletion of the last *Scn1a* exon, as well as in DS mice carrying the *Scn1a* A1783V missense mutations; both on the pure C57BL/6J background (Fadila et al., 2020; Beretta et al., 2022) (Table 1).

In conclusion, the *Scn1a*^WT/R613X^ DS model, harboring the recurrent nonsense *Scn1a* mutation and available in open access distribution, including to for-profit organizations through The Jackson Laboratory (Strain # 003771), demonstrates multiple Dravet-associated phenotypes with robust and specific deficits that can be used as a preclinical model for drug development. Thus, as current treatment options in Dravet are limited and considerable efforts are being made to produce novel and more effective anti-seizure small molecule drugs, as well as disease-modifying genetic treatments (Isom and Knupp, 2021), including those specifically targeting nonsense mutations, we propose that this model can provide a useful and powerful pre-clinical platform for the Dravet research scientific community.

## Author contribution statement

This work was performed in partial fulfillment of the requirements for a Ph.D. degree of A.M. A.M, E.C, J.A.A and M.R conceived the project. A.M, M.B, V.W, H.B, Y-H.C, V.S.D, and M.R designed, carried out the experiments and analyzed the data. All authors contributed to the article and approved the submitted version. We wish to thank all the members of the Rubinstein lab for their constructive criticism and technical support, as well as Natasha Stark (Tevard Biosciences) for technical support.

## Author discloser

J.L, V.W, H.B, Y-H.C, and V.S.D are current employees of Tevard Biosciences. E.C, V.S, and J.A.A are/were employed by the Dravet Syndrome Foundation Spain. A.M, M.B and M.R have no conflicts of interest to disclose.

