## Supplementary material for "Heat-induced seizures, premature mortality, and hyperactivity in a novel *Scn1a* nonsense model for Dravet syndrome": Table 1

|  |  | ***Scn1a* microdeletions** | | | | | | ***Scn1a* nonsense mutations** | | | | ***Scn1a* missense mutations** | | | | ***Scn1a poison exon*** | |
| --- | --- | --- | --- | --- | --- | --- | --- | --- | --- | --- | --- | --- | --- | --- | --- | --- | --- |
| **DS-models** | Δ exon 26  (Yu et al., 2006) | Δ exon 1  (Miller et al., 2014) | Δ exon 1 138 nt.  (Morey et al., 2022) | Δ exon 25  (Cheah et al., 2012)    (conditional) | Δ exon 8  (Jansen et al., 2020)  (conditional) | Δ exons 8-12  (Uchino et al., 2021) | Δ exon 7  (Ogiwara et al., 2013)  (conditional) | R1407X  (Ogiwara et al., 2007) | E1099X  (Tsai et al., 2015) | Exon 6 stop  (Valassina et al., 2022) | R613X | | H939R  (Dyment et al., 2020) | R1648H with induction of hyperthermic seizures  (Martin et al., 2010) | A1783V  (Kuo et al., 2019; Ricobaraza et al., 2019)  (conditional) | | Exon 20N  (Voskobiynyk et al., 2021) |
| **Genetic background for DS mice** | C57BL/6J | 50:50 C57BL/6J:129S6/SvEvTac | 50:50 C57BL/6J:129S6/SvEvTacAusb | C57BL/6J | C57BL/6J | C57BL/6J | C57BL/6J | C57BL/6J | 75% C57BL/6JNarl | C57BL/6J | 50:50 C57BL/6J:129S1/SvImJ | | C57BL/6NCrl | 50:50 C57BL/6J:129P2/OlaHsd | C57BL/6J | | C57BL/6J |
| **Heat-induced seizures <P18** | Not susceptible  (Oakley et al., 2009) | 40 - 42 °C  (Hawkins et al., 2016, 2017; Satpute Janve et al., 2021; Anderson et al., 2022; Gerbatin et al., 2022a, 2022b) | NA | NA | NA | NA | Mice with a global expression of the mutation were not studies | NA | NA | NA | Fig. 3  40 ± 0.27 °C | | NA | NA | ~ 38.5 -41 °C -  (Kuo et al., 2019; Almog et al., 2021) | | NA |
| **Heat-induced seizures P18-P28** | 38 - 41 °C  (Oakley et al., 2009; Rubinstein et al., 2015) | 38.5- 42 °C  (Nomura et al., 2019; Niibori et al., 2020; Hawkins et al., 2021a; Gerbatin et al., 2022a; Tanenhaus et al., 2022) | NA | ~ 38 °C  (Williams et al., 2019; SH et al., 2021) | NA | NA | NA | 40 - 41 °C  (Tanenhaus et al.; Cao et al., 2012; Gheyara et al., 2014) | ~ 40 °C  (Tsai et al., 2015) | NA | Fig. 3  38.78 ± 0.1 °C | | Increase number of seizures at 30 °C ambient temp of  (Dyment et al., 2020) | ~41.5 °C  (Salgueiro-Pereira et al., 2019) | ~ 37 – 39.5 °C  (Almog et al., 2021; Miljanovic et al., 2021) | | NA |
| **Heat-induced seizures >P30** | 38 - 39 °C  (Oakley et al., 2009; Cheah et al., 2021) | 40 – 42 °C  (Tran et al., 2020; Mattis et al., 2022) | NA | NA | 38.5 when crossed with the global EIIA-Cre (Jansen et al., 2020) | NA | NA | 40 - 42 °C  (Cao et al., 2012; Yamagata et al., 2020; Hawkins et al., 2021a) | ~ 40 °C  (Tsai et al., 2015; Hsiao et al., 2016; Ho et al., 2021) | ~ 40.5 °C  (Valassina et al., 2022) | Fig. 3  39.64 ± 0.46 °C | | NA | NA | 38 - 40 °C (Ricobaraza et al., 2019; Almog et al., 2021; Mora-Jimenez et al., 2021; Pernici et al., 2021) | | NA |
| **Hyperactivity in the open field** | +  (Han et al., 2012) | +  (Niibori et al., 2020; Hawkins et al., 2021b; Gerbatin et al., 2022a) | +  (Morey et al., 2022) | +  (Williams et al., 2019) | NA | NA | NA | +  (Ito et al., 2013; Gheyara et al., 2014; Yamagata et al., 2020) | NA | +  (Valassina et al., 2022) | +  Fig. 4 | | NA | +  (Salgueiro-Pereira et al., 2019) | +  (Ricobaraza et al., 2019; Fadila et al., 2020; Miljanovic et al., 2021; Mora-Jimenez et al., 2021; Satta et al., 2021) | | +  (Voskobiynyk et al., 2021) |
| **Increased anxiety** | +  (Han et al., 2012) | +  (Bahceci et al., 2020; Patra et al., 2020) | No signs of increased anxiety  (Morey et al., 2022) | +  (Williams et al., 2019) | NA | NA | NA | +  (Ito et al., 2013; Yamagata et al., 2020) | NA | No signs of increased anxiety  (Valassina et al., 2022) | No signs of increased anxiety  (Fig. 4) | | NA | No signs of increased anxiety  (Salgueiro-Pereira et al., 2019) | +  (Ricobaraza et al., 2019; Fadila et al., 2020; Mora-Jimenez et al., 2021) | | NA |
| **Motor deficits** | +  (Beretta et al., 2022) | NA | NA | NA | NA | NA | NA | NA | NA | NA | Normal performance of the rotarod  (Fig. 4) | | NA | NA | +  (Ricobaraza et al., 2019; Fadila et al., 2020; Mora-Jimenez et al., 2021; Alonso et al., 2022) | | NA |
| **Cognitive deficits** | +  (Han et al., 2012; Beretta et al., 2022) | +  (Bahceci et al. 2020) (Patra et al., 2020) | NA | +  (Williams et al., 2019)) | NA | NA | NA | +  (Ito et al., 2013; Gheyara et al., 2014) | NA | +  (Valassina et al., 2022) | Normal performance in the Y maze test  (Fig. 4) | | NA | +  (Salgueiro-Pereira et al., 2019) | +  (Ricobaraza et al., 2019; Fadila et al., 2020; Mora-Jimenez et al., 2021; Alonso et al., 2022) | | Not observed  (Voskobiynyk et al., 2021) |
| **Autistic features** | +  (Han et al., 2012; Beretta et al., 2022) | +  (Bahceci et al., 2020; Patra et al., 2020; Gerbatin et al., 2022a) | NA | +  (Williams et al., 2019) | NA | NA | NA | +  (Ito et al., 2013; Gheyara et al., 2014; Yamagata et al., 2020; Shao et al., 2022) | NA | +  (Valassina et al., 2022) | NA | | NA | +  (Salgueiro-Pereira et al., 2019) | +  (Miljanovic et al., 2021; Satta et al., 2021; Alonso et al., 2022) | | Not observed  (Voskobiynyk et al., 2021) |
| **Ordering information** |  | MMRRC / JAX  Strain #037107-JAX  Commercial License Agreement is required for for-profit use. |  | MMRRC  041829-UCD  Non-profit institutions only. |  |  | RBRC  MGI:5523787  MTA is required | RBRC  RBRC09420  MTA is required |  |  | JAX  Strain #:034129 | |  |  | JAX  Strain #:026133 | |  |
